# Predicting brain age across the adult lifespan with spontaneous oscillations and functional coupling in resting brain networks captured with magnetoencephalography

**DOI:** 10.1101/2024.01.10.574995

**Authors:** Samuel Hardy, Gill Roberts, Matthew Ventresca, Benjamin T Dunkley

## Abstract

The functional repertoire of the human brain changes dramatically throughout the developmental trajectories of early life and even all the way throughout the adult lifespan into older age. Capturing this arc is important to understand healthy brain ageing, and conversely, how injury and diseased states can lead to accelerated brain ageing. Regression modelling using lifespan imaging data can reliably predict an individual’s brain age based on expected arcs of ageing. One feature of brain function that is important in this respect, and understudied to date, is neural oscillations - the rhythmic fluctuations of brain activity that index neural cell assemblies and their functioning, as well as coordinating information flow around networks. Here, we analysed resting-state magnetoencephalography (MEG) recordings from 367 healthy participants aged 18 to 83, using two distinct statistical approaches to link neural oscillations & functional coupling with that of healthy ageing. Spectral power and leakage-corrected amplitude envelope correlations were calculated for each canonical frequency band from delta through gamma ranges. Spatially and spectrally consistent associations between healthy ageing and neurophysiological features were found across the applied methods, showing differential effects on neural oscillations, with decreasing amplitude of low frequencies throughout the adult lifespan, and increasing high frequency amplitude. Functional connectivity within and between resting-state brain networks mediated by alpha coupling generally decreased throughout adulthood and increased in the beta band. Predictive modelling of brain age via regression showed an age dependent prediction bias resulting in overestimating the age of younger people (<40 years old) and underestimating the age of older individuals. These findings evidence strong age-related neurophysiological changes in oscillatory activity and functional networks of the brain as measured by resting-state MEG and that cortical oscillations are moderately reliable markers for predictive modelling. For researchers in the field of predictive brain age modelling with neurophysiological data, we recommend attention is paid to predictive biases for younger and older age ranges and consider using specific models for different age brackets. Nevertheless, these results suggest brain age prediction from MEG data can be used to model arcs of ageing throughout the adult lifespan and predict accelerated ageing in pathological brain states.

## Introduction

Brain activity changes throughout the lifespan, rapidly in the early life developmental arcs during brain maturation (Hunt et al., 2019; Manza et al., 2015), through early-middle adulthood, and into older age (Vlahou et al., 2014). Neural oscillations offer a window in brain development, maturation, and ageing (Gómez et al., 2013), capturing neurophysiological processes at the population level (Rempe et al., 2023). Oscillatory activity can be captured via electroencephalography (Srinivasan et al., 2006) and magnetoencephalography (MEG). MEG is a functional neuroimaging technique that records the neuromagnetic activity of the brain via an array of magnetometers/gradiometers. Resting-state MEG is used to assess the spontaneous and intrinsic neural activity of individuals not engaged in an explicit task, revealing neurophysiological activity with minimal cognitive load. The benefit of MEG compared to other functional neuroimaging modalities such as fMRI and EEG is the greater temporal and spatial resolution, respectively, facilitating collection of ‘richer’ datasets in equal time, while remaining completely non-invasive despite directly recording neural activity. Resting-state MEG has been shown to be sensitive to age-related changes in neurophysiological activity (Hunt et al., 2019; Rempe et al., 2023; Vlahou et al., 2014), as well as a biomarker source for a diverse set of neurological and psychiatric states (O’Reilly et al., 2017; Huang et al., 2014; Dunkley et al., 2014; Allen et al., 2021), and can even differentiate between disorders with significant symptom overlap (Zhang et al., 2021).

Age is a primary risk factor for most neurodegenerative diseases, and MEG has been leveraged in studies of neurodegeneration such as Alzheimer’s and other dementias (Wiesman et al., 2022; Engels et al., 2016) and Parkinson’s (Vardy et al., 2011). Characterising the typical course of healthy ageing in the adult human brain will help us to understand the importance of deviation from expected trajectories of age dependent neurophysiological features, and if such deviation is associated with neurodegeneration and accelerated neural ageing (Ye et al., 2020). Functional neuroimaging derived features would define new pathways for the early detection of pathological brain states, beneficial in cases where pre-symptomatic pathology is most effectively treated with early intervention.

Neural oscillations are a key feature of neural processing and reflect rhythmic activity of neural assemblies, across multiple spatial and temporal scales. Importantly, they have been implicated in information processing and transfer, perceptual binding, cognitive processes, and behavioural output. Furthermore, they provide mechanistic information about disease state (Mathalon & Sohal, 2015). Oscillatory activity can be recovered from neural spectral power, and is a biophysical phenomenon used to describe the activity of frequency specific processing throughout the brain. Associations between healthy ageing and M/EEG detected neural activity generally detail a profile of consistent spectral changes, including oscillatory activity shifting from low to high frequencies, as measured via band limited power (Barry & De Blasio, 2017) and full power spectra (Voytek et al., 2015; Merkin et al., 2023). These spectral changes are not spatially uniform across the cerebral cortex, evidenced by theta and alpha showing the strongest decreases in the posterior regions, and conversely beta activity being most acutely increased across the frontal hemispheres (Barry & De Blasio, 2017).

Neural oscillations also mediate functional connectivity (FC), and in electrophysiological contexts, FC is usually defined by phase-or amplitude-based coupling modes (Engel et al., 2013). Amplitude envelope correlations (AEC) however, are generally the most stable (Colclough et al., 2016) and reliable over time (Garcés et al., 2016), and intrinsic resting networks recovered from AEC closely resemble those recovered by other imaging modalities, such as more widely used functional MRI (fMRI) (Brookes et al., 2011a). Studies of resting-state FC using fMRI have demonstrated age related changes in the amygdala (Xiao et al., 2018) and basal ganglia (Manza et al., 2015) between young and middle adulthood, as well as in the default mode network (Ferreira & Busatto, 2013). While fMRI measures neural activity indirectly via blood-oxygen-level-dependent (BOLD) signals with a temporal resolution of approximately 1-5 seconds, MEG directly measures the neuromagnetic activity of the brain with millisecond resolution, allowing connectivity to be measured without the limitation of the haemodynamic response time. Nevertheless, there is excellent spatial and spectral congruence between MEG and fMRI measures of resting-state FC (Brookes et al., 2011a; Brookes et al., 2011b), and MEG-based studies have revealed differences in network topology between children, adolescents and early-middle adulthood (Schäfer et al, 2014).

The relationships between neural activity and age are often improperly modelled as purely linear, however the true trajectories of these features with age appear to be generally monotonic, though quadratic relationships are also present (Gómez et al., 2013; Hunt et al., 2019) but are weak enough to have most variation can be captured by a monotonic fit. Additionally, research in this area is often limited by mixed modality use (Voytek et al., 2015), limited sample sizes (Voytek et al., 2015; Barry & De Blasio, 2017), narrow age ranges (Schäfer et al, 2014; Hunt et al., 2019), and group-wise rather than continuous analysis of ageing (Merkin et al., 2023; Koelewijn et al., 2017), which highlights the need for further analysis of large M/EEG datasets in this area which comprehensively cover the typical adult human lifespan in a continuous manner.

Using neuroimaging modalities to predict age is an established technique, commonly referred to as “brain age” (Cole et al., 2019). This process involves modelling the effect of age on neuroimaging data as collected from many samples to account for personal variation and limit bias. As variation is present in functional and structural brain data within even limited age groups, predictions (i.e. brain age) frequently differ from chronological age, with the difference being called residuals or brain age delta (BD). Since BD is found by subtracting the real value of age from brain age, a positive BD means a predicted age higher than chronological age, which is a result of brain data which more closely resembles data from an individual older than chronological age would imply. Given the potential for difference between chronological and brain age, research has been conducted to explore if this residual error is related to biomarkers of various pathologies, primarily but not limited to those of a neurological nature (Lee et al., 2022; Rauseo et al., 2023; Ye et al., 2020).

This work will explore age-related effects on neurophysiological features including spontaneous cortical activity and FC derived from resting-state MEG data. We replicate prior findings on cortical oscillatory and network activity, and advance the field significantly by revealing the age-related changes from early-to-late adulthood in intrinsic functional networks, a novel approach with MEG data. In doing this we also determined stable features of maturation across the adult lifespan, allowing comparison to existing literature in this subject area. Additionally, we performed regression analysis to predict chronological age using MEG derived biophysical features of brain activity, and show that brain age predictive modelling using cortical oscillatory activity on the whole is moderately reliable, but that there are small age-related biases in predictions that researchers using neurophysiological data should be mindful of.

## Methods

### Participants

Participants data was compiled from four distinct cohorts and sites, including The Hospital for Sick Children Toronto (n=18), The Open MEG Archive (OMEGA) healthy controls (n=106), National Institute of Mental Health (NIMH) healthy research volunteers (n=49), and a subset of the Donders Institute Mother of Unification Studies (MOUS) (n=194). For each participant, we utilised T1 MRI for source-localisation and eyes-open resting-state MEG neuroimaging data. The total cohort size is 367 participants (mean age=36.51±20.07 years, age range=18-83 years, female=59.94%). All data were collected under ethical approval from each respective site’s Research Ethics Board / Institutional Review Board (REB/IRB), and all participants provided informed consent to participate in research.

### MEG Data & Processing

#### Pre-processing

All MEG data were processed with a custom pipeline utilising the open source MNE-Python software package (Gramfort et al. 2013). Recordings were temporally cropped to the first 5 minutes of the recording, then were band pass filtered between 1 and 150 Hz and resampled to a frequency of 300 Hz before processing. A notch filter was used to remove line noise at 50 and 60 Hz.

CTF and MEGIN systems have different sensor arrays, with separate magnetic interference rejection mechanisms. For data acquired with CTF systems, synthetic 3rd-order gradiometry using a reference array was used to attenuate interference. For data acquired using MEGIN systems, signal-space projections were computed using both magnetometers and gradiometers and then applied to the filtered data. Subsequently, independent components analysis was performed with 80 components identified for each scan using the infomax approach (Lee et al. 1999). Artefactual components (ocular, cardiac) were identified automatically using a proprietary machine learning classifier and excluded.

Lastly, the sensor-level data were segmented into 10.24-second epochs. Any epochs with a peak-to-peak amplitude greater than 6000 fT were discarded from further analysis; any epochs with high-frequency (>110 Hz) activity greater than 2000 fT were also discarded, a threshold which was chosen to reject epochs containing high-frequency muscle activity. Remaining clean epochs were discarded until 250 seconds of data were available, ensuring that the same length of data was analysed for all scans.

#### Forward modelling

We used a 1-shell boundary element model, with the geometry warped to fit each subject’s anatomical MRI, to calculate magnetic lead fields (Nolte 2003); these lead fields were subsequently used in the inverse solution for neural source reconstruction. Specifically, the subject’s T1 MRI was resampled to isotropic 1 mm resolution and underwent an affine (non-rigid) registration to a template brain using Freesurfer software (Fischl 2012). This registration defines a correspondence between individual subject source spaces and the template space. The template includes precomputed boundary meshes defining skull compartments; we transformed the inner skull mesh, along with source locations of interest, back to each subject-specific source space.

Dipolar sources were modelled at 122 locations across the brain. These fall into two categories: 78 locations from the AAL atlas (Gong et al., 2009) which were used for regional power estimates, and 44 other MNI coordinates which were used as network nodes (**Appendix 1**). These locations were selected a-priori to model five specific large-scale networks: the central executive (CEN), default mode (DMN), motor (MOT), visual (VIS), and attention (ATT) networks.

The specific MNI coordinates were adapted from previous work in the field (De Pasquale et al., 2012; Gong et al., 2009; Hsu et al. 2014); specific node groupings may be found in **Appendix 1**.

#### Source imaging

For each subject, cleaned epochs were used to compute a covariance matrix for source reconstruction using a LCMV beamformer (Vrba and Robinson, 2001). We applied regularisation to the covariance matrix to increase numerical stability of the inversion, using 5% of the maximum singular value applied to the diagonal elements of the matrix. Epochs were spatially filtered at the 122 source locations using three independent cartesian components for each source; these were reduced to scalar source time courses using a singular value decomposition method (Sekihara 2004).

We estimated power spectral density for source time courses at each of the 78 AAL locations using Welch’s method. The resulting power spectra were divided into five a-priori frequency bands and the mean amplitude for each frequency band was computed. This resulted in estimates of band-limited spectral power at each AAL node.

Source time courses from the network nodes were used to estimate connectivity. Firstly, the time courses were filtered in each frequency band, then orthogonalized using a symmetric procedure to remove zero-lag correlations; this removes any spurious correlations which would otherwise result from magnetic field spread (Colclough et al. 2015). We then used the Hilbert transform to estimate an amplitude envelope for each source time course. These envelopes were downsampled to 1 Hz and the Pearson correlation coefficient between pairs of nodes was calculated. The output takes the form of a NxN matrix, where each element defines the correlation between a pair of nodes.

#### Partial Least Squares

Partial least squares (PLS) is a cross decomposition method similar to principal component analysis (PCA), but whereas PCA reduces the dimensionality of a matrix such that the resulting components maximally explain the variance of the matrix (while remaining orthogonal), PLS instead produces two sets of components, between pairs of which the covariance is maximised (Abdi & Williams, 2013). PLS is useful in high dimensional data in which dimensionality reduction with respect to a specific set of targets is ideal, and further where the regressors may be multi-collinear. We used PLS to explore the relationships between MEG data and metadata such as age and sex, as a means of further exploring how these variables are represented neurophysiologically. To perform PLS we begin with the calculation of the cross-product matrix as shown in equation 1, in which **X** is the independent variables (MEG features), and **Y** is the dependent variables (metadata), where **X** and **Y** have both been centred and normalised.

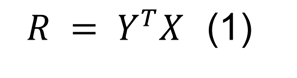

The singular value decomposition (SVD) of **R** is then calculated using equation 2, where **U** and **V** are the left and right singular vectors respectively, and **Δ** is the singular values. Right and left singular vectors represent the symptoms and MEG features respectively which best describe **R.**.

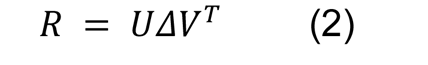

Components (also called latent variables) are then produced via projection of the MEG features onto the left singular vector (**V**), and the symptom scores onto the right singular vector (**U**) as shown in equations 3 and 4.

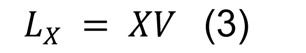

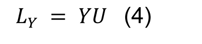

These latent variables are often called scores, and pairs of scores such as **Lx0** and **Ly0** are designed to be maximally covariant and represent the relationship between the **X** and **Y** features to the extent that the common information in the two matrices allows.

To test the significance of the produced latent variables we compare the singular values (**Δ**) of the SVD of **X** and **Y** to null distributions of the singular values generated by permutation testing. This process involves shuffling **X** and recomputing the singular values via a new SVD under the assumption of no relationship between **Y** and shuffled **X**. For each singular value of the original SVD we can use the corresponding null distribution to calculate the p-value which describes the probability of witnessing a singular value at least as extreme as observed. It is possible for only a subset of the singular values of an SVD to be found statistically significant (α=0.05), and in this case only those which are significant are considered for further analysis.

In assessing the relationship between the latent variables of **Y** (**Ly**) and the features (columns) of **Y** we first used a spearman’s rank correlation. Furthermore, we produced confidence intervals of these correlations via bootstrapping, whereby 1000 resamples with replacement are performed, and the SVD and subsequently the latent variables are computed for each, with the correlation coefficients between these latent variables and the bootstrapped **Y** features being used to generate confidence intervals.

#### Regression

For all modelling the regressor variables of MEG derived spectral power and FC features are used to predict the continuous response variable of chronological age.

Sci-kit Learn (Pedregosa et al., 2011) is used for its cross-validation, standardisation, and modelling functionality in this study. To evaluate the performance of each model we used repeated K-Fold (k=10) cross-validation, in which 10 repetitions are carried out. Additionally, we stratify the response variable of age into 4 linearly spaced groups (using the limits 0.0, 22.5, 45.0, 67.5, 90.0), to ensure that each fold contained an adequate age distribution. Response variable stratification is used to minimise the effect of skewed response variable distributions in training folds limiting the generalisation of models. Although stratification is performed over grouped ages, prediction is performed against the continuous ungrouped value of chronological age. The performance of all models is measured using mean absolute error and r-squared, with the mean and standard deviation of these metrics for training and testing data over cross-validation being recorded.

To optimise the parameters of each of the tested regression models we performed a cross validated parameter grid search over a range of suitable parameter values, using the same cross validation procedure previously specified. For each model architecture we select the best performing parameterisation to compare against alternative architectures.

Existing literature details at length a common pitfall of predicting age using neuroimaging features as prediction bias, particularly over-predicting (positive residuals) for younger subjects, and under-predicting (negative residuals) for older subjects. Multiple bias correction methods exist which attempt to address this issue (Peng et al., 2021; de Lange et al., 2020), and following the reasoning specified we chose to train a regression model (on the train folds) using chronological age to predict the age prediction of the main model, the training and testing data is then bias corrected by subtracting the intercept from the main model predictions and dividing by the gradient as shown in equations 5 and 6, where **x**, **y**, and **x’** are brain age, chronological age, and bias corrected brain age respectively.

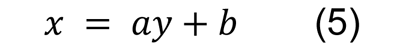

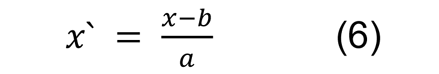

This approach ensured no data-leakage in the bias-correction process; however, it does assume that the bias of the training data predictions is approximately equal to that of the test data predictions. Given that the difference in the bias between training and testing predictions is a proxy for over/underfitting, it is necessary to minimise the difference between training and testing performance metrics while optimising these metrics, i.e. minimising mean absolute error and maximising r-squared. To mitigate the effect of this disparity in bias between train and test sets we also test using a subset of the training fold in each cross-validation iteration as a specific bias correction holdout set, which due to not being used for model training, would have approximately equal bias to the test set, this holdout set could therefore be used to train bias correction linear model.

## Results

### PLS reveals cortical oscillatory and connectomic changes associated with healthy ageing

Cortical oscillatory activity shows strong age dependent effects, with low frequencies, including delta and theta, showing global decreases across the adult life span (Figure 2a, 2b), while high frequencies, such as beta and gamma, show widespread increases (**Figure 2d, 2e**), with alpha showing spatially specific differential relations. Additionally, the coefficients of spectral power shown in Figure 2 present the same general trends with respect to age as the partial correlation coefficients (**Appendix 2,** **Figure S1**), specifically, decreasing delta activity except in the occipital lobe, decreasing theta focussed in the temporal and parietal regions, decreasing occipital alpha and increasing temporal alpha, increasing global beta power, and increasing gamma across the whole brain but prominently in frontal areas and most weakly in temporal regions. Generally, the effect of healthy ageing found here is a shift toward high frequency activity, with the alpha band capturing the pivot point around which activity shifts, evidenced by its status as the only frequency band exhibiting bi-directional changes associated with healthy ageing.

**Figure 1:**
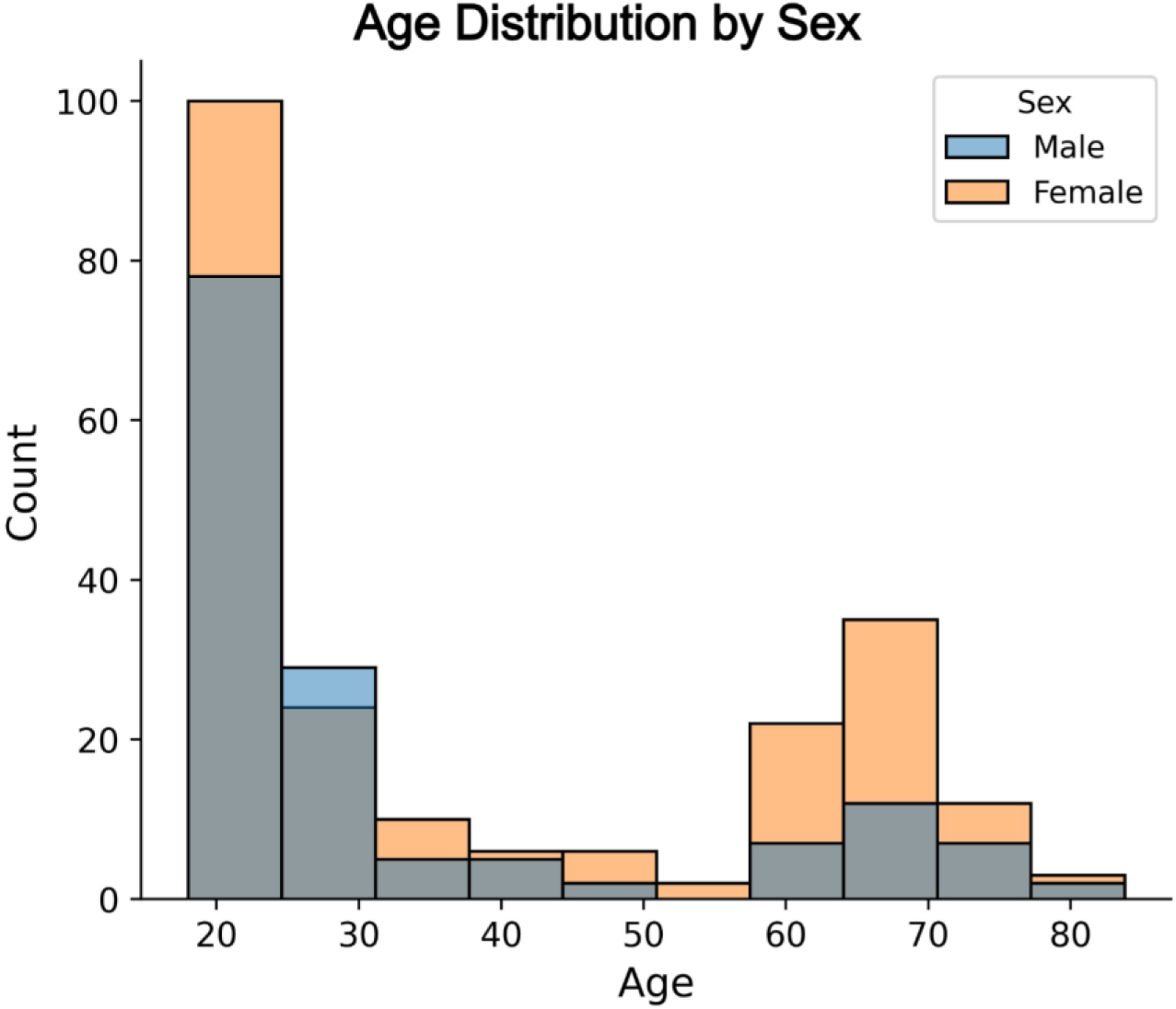
Histogram of participant age, grouped by sex.

**Figure 2:**
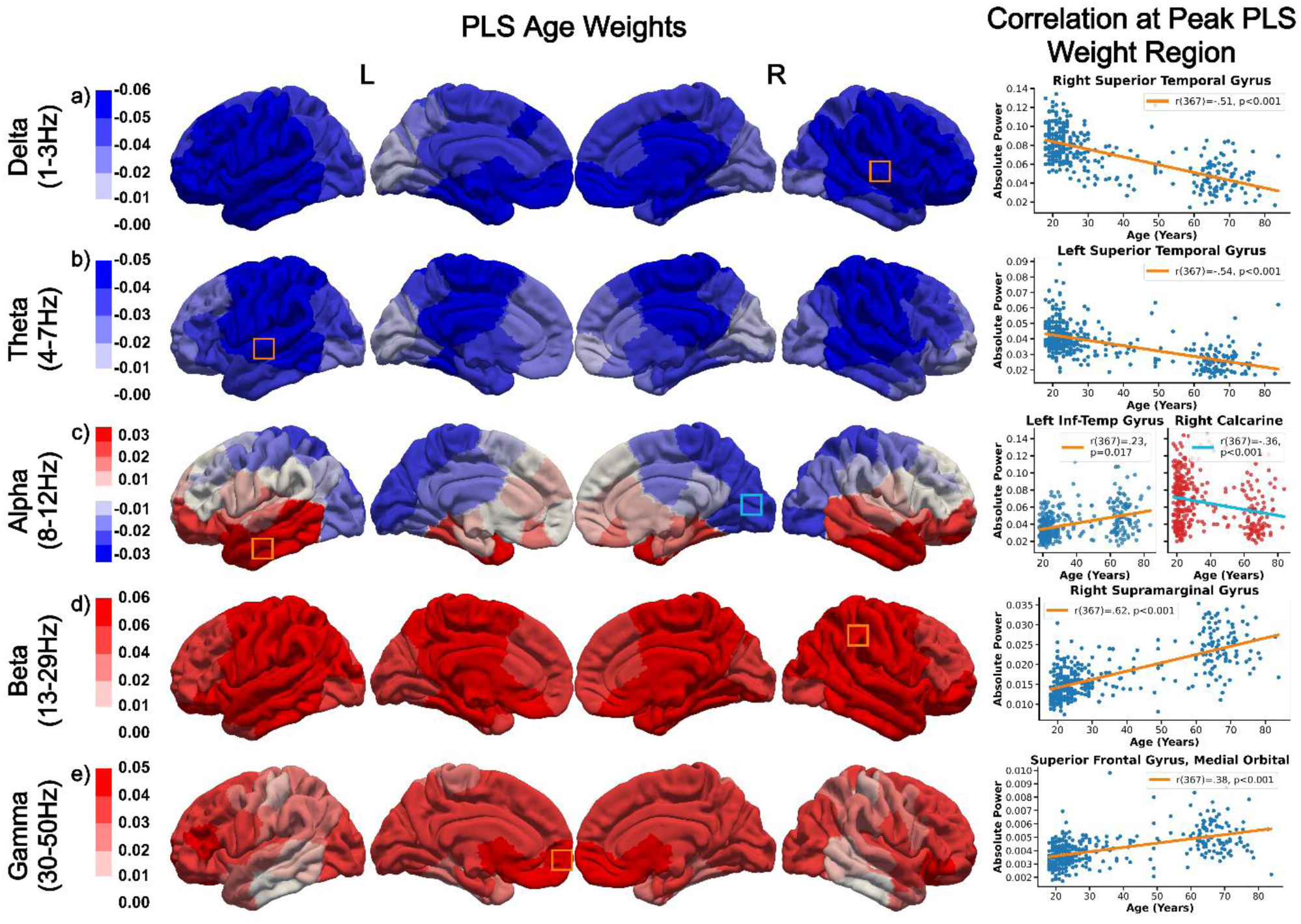
Spectrally and spatially specific effects of ageing on cortical oscillatory activity. Low frequency (e.g. delta and theta) decreases in power with age are prominent in frontal and temporal regions, with decreases in alpha seen in the occipital regions. Age associations in alpha are diminished with respect to the spectrally adjacent theta and beta bands, which exhibit stronger and exclusively unidirectional changes. The most pronounced age-related changes in power are in beta, in which increasing activity across all brain regions is associated with healthy ageing. Similarly, gamma activity was found to diffusely increase with ageing, primarily in frontal regions. Bounding boxes on the band limited power plots highlight PLS coefficient peak regions, the colour of the bounding box corresponds to the line of best fit shown on the adjacent scatter plot, which shows data from this region.

The following regions have the largest frequency band specific absolute coefficient value in PLS analysis for predicting age, right superior temporal gyrus (***r***(367)=-.51, ***p***=6.24e-20), left superior temporal gyrus (***r***(367)=-.54, ***p***=1.30e-20), left inferior temporal gyrus (***r***(367)=-.36, ***p***=4.48e-9), right supramarginal gyrus (***r***(367)=.62, ***p***=1.43e-27), and left inferior temporal gyrus (***r***(367)=.38, ***p***=1.08e-9) respectively from the previously mentioned canonical frequency bands.

Healthy ageing was found to be associated with multiple network level features of FC, including decreases in the default mode network in delta (Figure 3a), decreases in the visual network in alpha, beta, and gamma (Figure 3c, 3d, 3e), as well as diffuse connectome wide increases in beta and gamma (Figure 3d, 3e). The average absolute PLS coefficient value of connectivity is significantly lower than that of the spectral power coefficients (***U***=107464, ***p***=9.33e-186), highlighting that spectral power features have greater association with age on average. This is also reflected in Figure 3 which shows the comparatively low coefficient values for connectivity. Furthermore, these results show region, frequency band, and directional concordance with partial correlation coefficients of the default mode of beta, as well as the visual network in alpha, beta, and gamma (**Appendix 2**, **Figure S2**).

**Figure 3:**
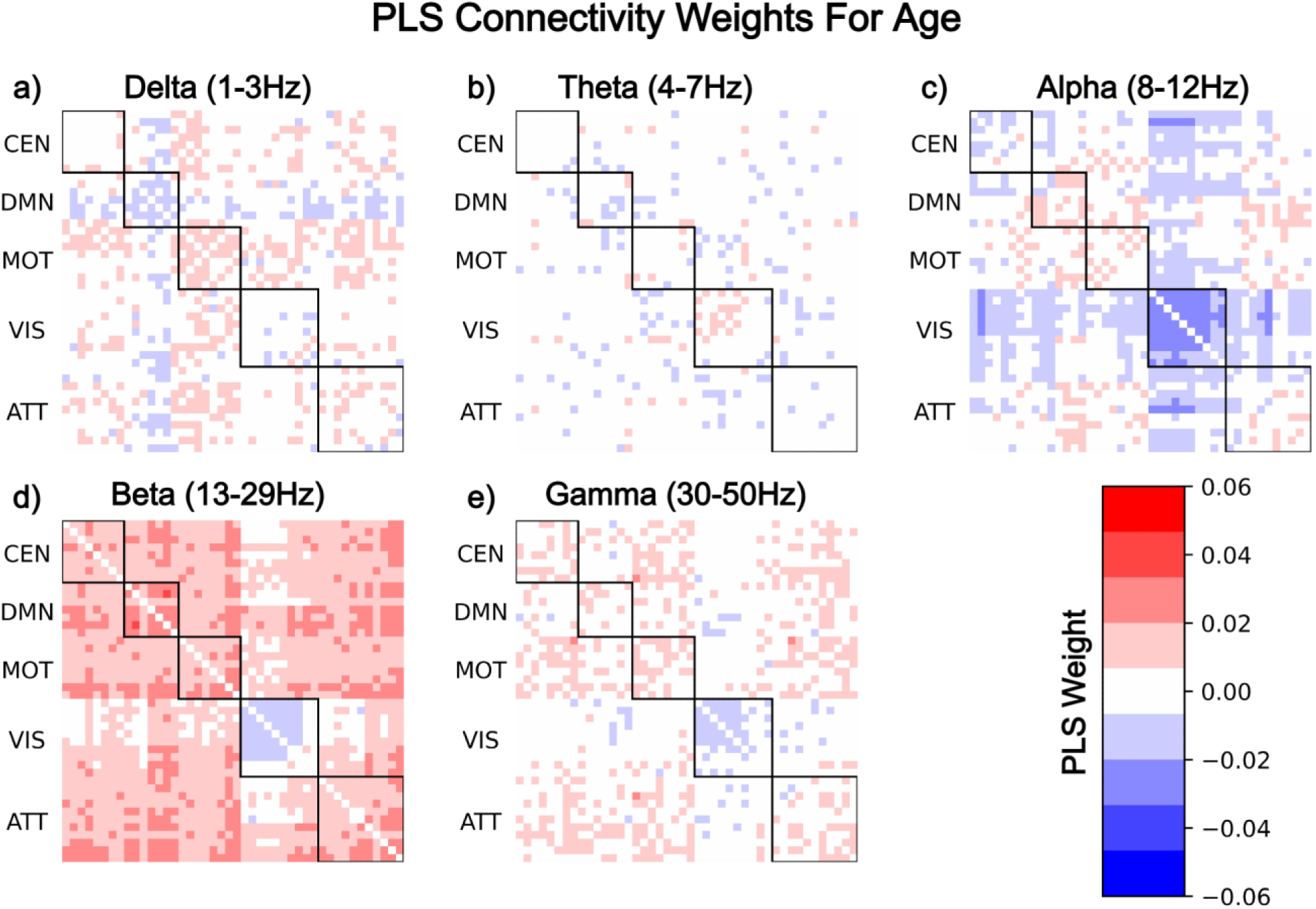
Functional connectivity shows network and frequency specific changes associated with healthy ageing, with inter and intra default mode network connectivity decreasing in delta, while theta exhibits no clear effect beyond single node connectivity changes. Connectivity is decreased within the visual network across alpha, beta, and gamma, with the effect most exaggerated in alpha, where inter visual network connectivity also decreases to a lesser extent. Outside of the visual network beta and gamma demonstrate diffuse increasing connectivity associated with healthy ageing.

#### MEG-derived oscillatory activity for predicting functional brain age

A variety of models provided in the sci-kit learn python package were evaluated via parameter optimisation in the model selection process, the best model which is subsequently used in the following sections is an adaboosted (Drucker, 1997) cross validated ridge regression model parameterised to have strong regularisation, and only use spectral power features. This particular model was selected as it produced the lowest summed value of mean test fold performance and the difference between train and test fold performances (as measured via mean absolute error). Using the difference between train and test performance metrics is intended to penalise overfit models which would not work well with the specified bias correction procedure. Various subsets of the features were used to test the models, as well as decomposition methods such as principal component analysis (PCA), with results showing that using PCA and only spectral power both independently reduced overfitting, though PCA also worsened test fold performance.

#### Raw Predictions

As expected, a degree of bias exists in the age predictions, as visible in Figure 4. It is also clear that the severity of the bias is increased in the test set compared to the training data, which, while anticipated, does limit the effectiveness of the bias corrected procedure later used, as the degree of difference is substantial. The prediction bias found follows the trend of existing literature in this domain, in which younger peoples ages are overpredicted (positive average brain age delta), and older individuals ages are underpredicted (negative average brain age delta). The use of age stratified cross-validation does appear to have limited the detrimental effects of the data sparsity for the middle-aged subjects, with test fold performance not suffering in this age range compared to others.

**Figure 4:**
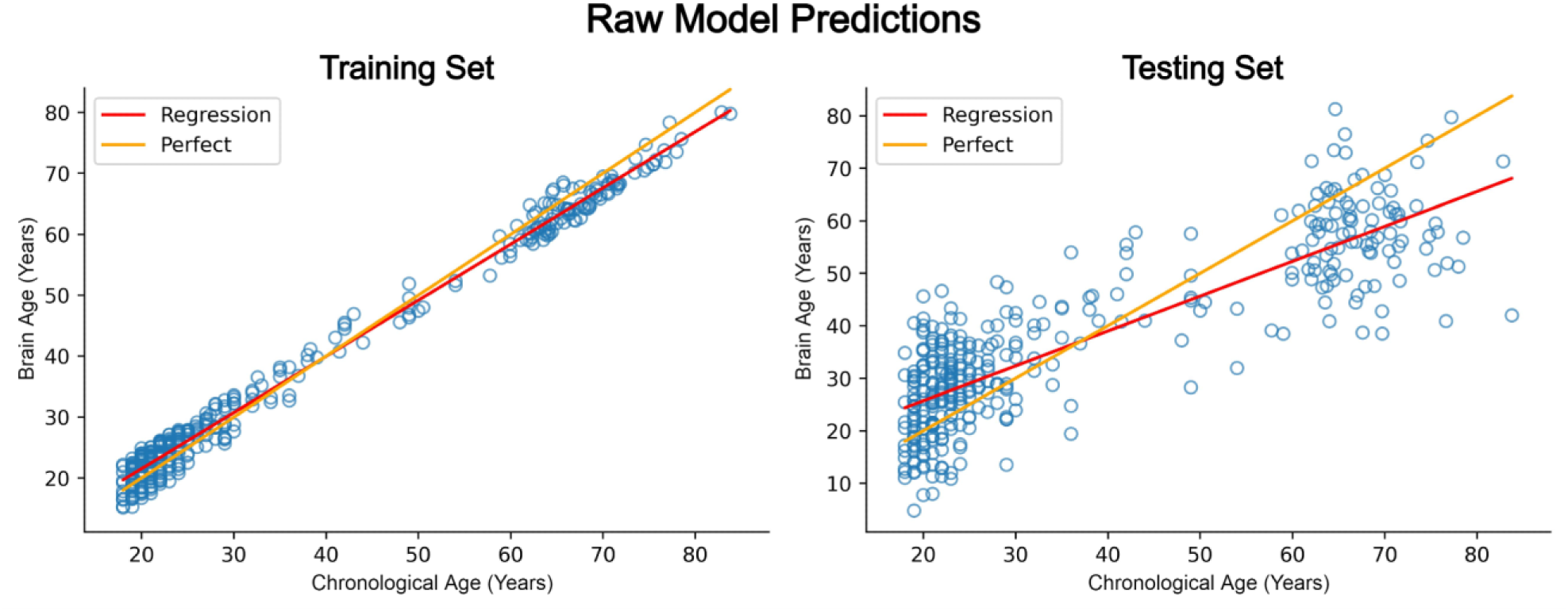
Model predictions show significantly reduced error over training samples compared to test set (unseen) samples, highlighting the presence of overfitting. Additionally, both training and testing predictions display age dependent prediction bias resulting in overestimating the age of younger people (<40 years old) and underestimating the age of older individuals (>40 years old), which is highlighted by the divergence of the ideal and actual lines of best fit of the predicted and actual values of age.

**Table 1:** The regression model demonstrates overfitting to the training samples, evidenced by the disparity between the performance measures of the train and test sets. The test samples are predicted with mean absolute error of 8.54 years.

|  | Mean Absolute Error | R<sup>2</sup> |
| --- | --- | --- |
| Train Fold Performance | 2.36 | .98 |
| Test Fold Performance | 8.54 | .71 |

#### Bias-Corrected Predictions

During each cross-validation iteration the training fold predictions and labels are used to calculate the bias correction linear model, this model is then used to adjust the test fold data, which are then stored in both raw and bias corrected form for each individual in the test fold. After all cross-validation iterations have completed the raw and bias corrected test fold predictions are averaged for each individual, creating the results shown in Figures 4 and **5**. As the bias correction models are calculated using the training folds to avoid data leakage, the bias of the test folds will only ever be adjusted to the degree that the bias is found in the train folds. If therefore the bias between these two data sets is substantially different, the bias of the test data will not be optimal, as seen in Figure 5, the right side of which highlights the amount of bias remaining (the degree of divergence between the regression and perfect fits) in the test data. Although in this case the bias correction is nonoptimal, it still demonstrates limited effectiveness as predictions are adjusted in the correct directions across the age continuum, but not to the ideal degree.

**Figure 5:**
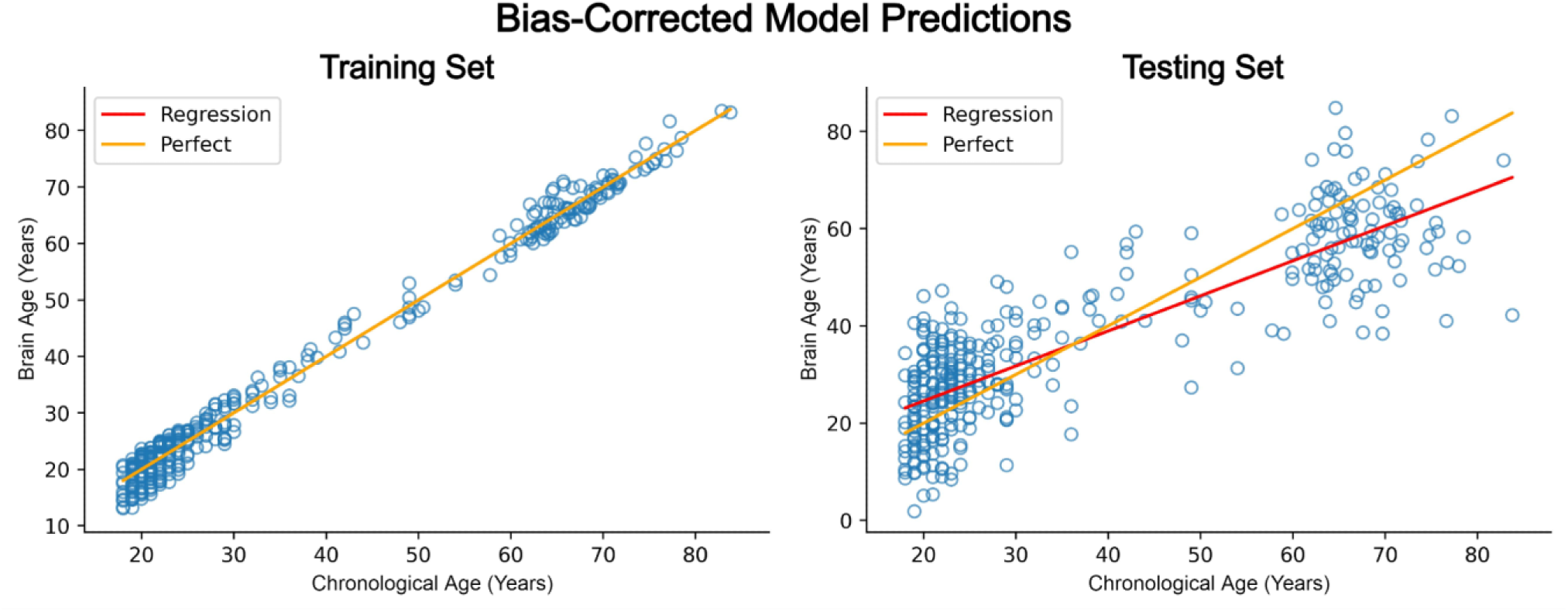
The effect of the bias-correction on the training set is clear, as the divergence between ideal and actual fits is removed, however the effect on the test samples is limited (due to overfitting), which is clear by the persistence of non-overlapping lines of best fit for ideal and actual test set predictions.

**Table 2:**
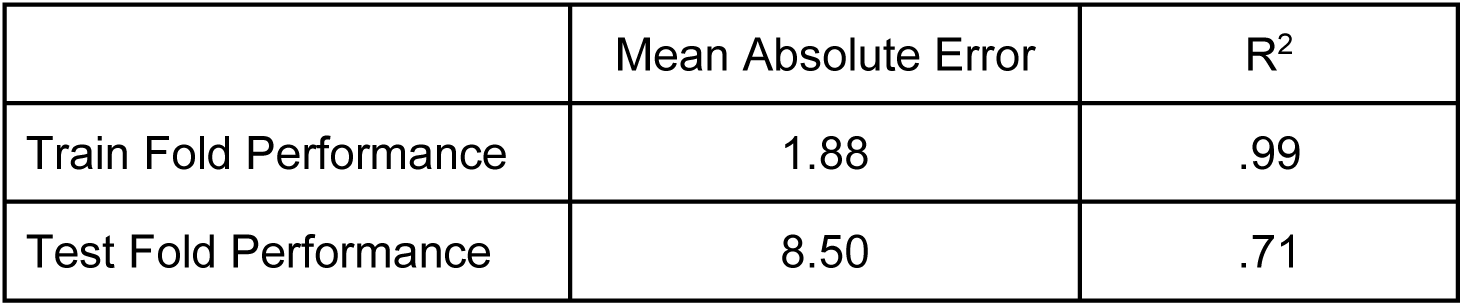
The performance values of the bias-corrected predictions are improved over the raw predictions, most prominently in the training performance, and to a lesser extent in the mean absolute error of test set performance.

|  | Mean Absolute Error | R<sup>2</sup> |
| --- | --- | --- |
| Train Fold Performance | 1.88 | .99 |
| Test Fold Performance | 8.50 | .71 |

## Discussion

### Summary

Neurophysiological activity develops and matures throughout the process of healthy ageing, reflecting underlying changes to cellular and population functioning. We sought to replicate prior work on the maturational effects of regional oscillatory activity, advance our understanding of oscillatory-mediated functional connectivity across resting-state brain networks, and test brain age prediction models on neurophysiological data. We replicated prior studies that found age-related decreases in low frequency activity (Barry & De Blasio, 2017; Hunt et al., 2019), while simultaneously high frequency activity (Barry & De Blasio, 2017; Gómez et al., 2013) tends to increase. Furthermore, we present novel results that show intrinsic coupling in key resting-state networks, including the DMN, CEN and attention network, including age-related decoupling mediated by alpha activity, especially within and between the visual network, and across network increases in coupling mediated by beta activity. Finally, we present a regression model that tested brain age prediction modelling on neurophysiological data and found that the age of younger individuals tends to be overpredicted and the age of older individuals is underpredicted, calling attention to the sensitivity of the tested bias-correction procedure to overfitting.

### Oscillatory activity throughout adulthood

Oscillatory power spectra tend to flatten during the course of healthy ageing (Voytek et al., 2015; Merkin et al., 2023), conversely, the effect of neurological diseases can cause a steeper spectra, due to increasing low frequency activity, and decreasing high frequency activity. These mutually occurring changes are characterised as neural slowing, in which the amplitude across global power spectra shifts toward lower frequencies. Multiple methods for modelling neural slowing have been proposed, including utilising the coefficients of 1/f (Voytek et al., 2015; Merkin et al., 2023) and mean frequency (Fernández et al., 2006) of power spectra, as well as linear models of band limited power z-scored using a normative group (Wiesman et al., 2022).

Irrespective of methodology, neural slowing decreases with healthy ageing (Voytek et al., 2015; Merkin et al., 2023). However, individual and group level deviations from the established spectral trajectory of age have been associated with the presence of neurodegenerative diseases. This has been established using both band limited power and spatially resolved neural slowing as a biomarker of AD (Wiesman et al., 2022; Giustiniani et al., 2023; Horvath et al., 2018; Mandal et al., 2018; Ishii et al., 2018; Maestú & Fernández, 2020; Chen et al., 2015) and Parkinson’s disease (Bočková & Rektor, 2019; Oswal et al., 2013; Vardy et al., 2011). Additionally, increasing delta activity has been shown to correlate with poorer cognitive performance in both healthy and mild cognitive impairment (MCI) participants (Cuesta et al., 2015), and decreasing alpha has been associated with worse cognitive scores along the control, MCI, mild AD continuum (Babiloni et al., 2006).

### Maturational effects on functional coupling in intrinsic brain networks

Ageing also causes FC to decrease in networks that contribute to higher cognitive functions such as the default mode, cingulo-opercular, and fronto-parietal control networks, as well as increase between networks such as the visual and somatomotor (Geerligs et al., 2015). Global FC has been shown to increase with age, prompted by increasing intra and inter network connectivity (Schäfer et al, 2014). Graph theoretic measures have also been applied to FC, with modularity and local efficiency being shown to decrease with respect to age (Geerligs et al., 2015). The phase locking value was found to be lower in alpha and beta frequency bands in MCI subjects compared to controls when at rest (López et al., 2014). Similarly, coherence and synchronisation likelihood are lower in MCI than in controls, with both measures being significant in the beta band (Gómez et al., 2009). A lower global FC has been witnessed in AD when compared to both subjective cognitive decline and control subjects, particularly in alpha and beta (Schoonhoven et al., 2022; Koelewijn et al., 2017).

### Predicting brain age from neurophysiological data

Multiple studies have assessed the applicability of neuroimaging in brain age prediction, with most literature utilising T1 MRI data to test and train regression models. Structural imaging techniques have largely shown greater predictive power across regression metrics for this task, in part due to the greater sample sizes, but primarily due to the presence of clear anatomical effects of ageing. Decreasing brain volume, white, and grey matter (Lockhart & DeCarli, 2014; MacDonald & Pike, 2021; Gunning-Dixon et al., 2009) alongside increasing ventricle and cerebral spinal fluid volume (MacDonald & Pike, 2021) give clear indications as to the age of the participant. Furthermore, associations between structural atrophy and cognitive decline are suggested, particularly with respect to cortical thinning rates (MacDonald & Pike, 2021) and white matter reductions (Gunning-Dixon et al., 2009). These trends further generalise to neurodegenerative diseases, in which similar regional dependent atrophy in grey and white matter is present (Young et al., 2020), and thus pathological neurodegeneration can appear in structural neuroimaging as an acceleration of the natural processes of ageing. This benefits the premise of predicted brain age as we naturally equate higher age to worse cognition.

Prediction of age via structural imaging of the brains of healthy subjects has been explored via a variety of algorithmic approaches (Xifra-Porxas et al., 2021; Cole et al., 2018; Peng et al., 2021; de Lange et al., 2022) which have achieved mean absolute error as low as 2.95 years (Peng et al., 2021). Given the success of this technique, research has also explored the relationship between brain age predictions/BD and various diseases. Brain age predictions on AD cohorts using structural imaging have shown higher BD compared to controls (Lee et al., 2022; Rokicki et al., 2021), as well as a progression of higher BD related to severity of neurodegeneration (Lee et al., 2022; Franke & Gaser, 2012) and cognitive impairment (Franke & Gaser, 2012; Poloni et al., 2022) within and across neurological diseases such as MCI, AD, preclinical AD, frontotemporal dementia, and dementia with Lewy bodies. Conversion between disease groups at a later stage has been related to BD at baseline (Lee et al., 2022; Franke & Gaser, 2012; Gaser et al., 2013). Higher BD has also been associated with ischemic heart disease (Rauseo et al., 2023), increased mortality risk (Cole et al., 2018), traumatic brain injury (Cole et al., 2015), and schizophrenia (Nenadić et al., 2017). However, it appears that higher BD does not generalise past schizophrenia to other psychiatric disorders (Nenadić et al., 2017) unless functional imaging is also utilised (Rokicki et al., 2021).

The benefit of multimodality in predicting brain age and relating BD to non-imaging data highlights that structural and functional imaging explain distinct features of importance with respect to neurophysiological disorders (Niu et al., 2020; Cole et al., 2020). Age prediction via functional imaging is possible due to the prior mentioned changes in neural oscillations and FC, namely the flattening of the power spectra alongside increasing intra/inter FC. Furthermore, functional imaging may be preferential for disorders which do not present with structural abnormalities such as atrophy. FC via functional magnetic resonance imaging (fMRI) has been used to predict brain age (Li et al., 2018; Millar et al., 2022) and has shown the same relationship between BD and the severity of neurodegeneration across preclinical to symptomatic AD cohorts (Millar et al., 2022). Electrophysiological neural dynamics of the brain have also been used to predict participant age via EEG and MEG, both at rest (Engemann et al., 2022; Sabbagh et al., 2019; Sabbagh et al., 2020; Vandenbosch et al., 2019; Al Zoubi et al., 2018) and during sleep (Sun et al., 2019; Ye et al., 2020; Paixao et al., 2020), with further replication of increased BD being linked to life expectancy, neurological and psychiatric disorders between clinical groups as compared to healthy controls (Sun et al., 2019; Ye et al., 2020; Paixao et al., 2020).

Age dependent prediction bias is present in our regression results, with positive and negative BD for younger and older participants respectively. This phenomenon is present throughout the related literature (More et al., 2023; Smith et al., 2019; Peng et al., 2021) and we evaluated a method of bias correction explored in other studies, though this resulted in minimal bias reduction due to a large difference in fit between train and testing data (i.e. overfitting). Despite the limited effectiveness of the chosen correction method in this analysis, the application of bias correction did result in a decrease in mean absolute error for both train and test sets, highlighting the benefit of such techniques in this domain. However, correlation of bias corrected BD to non-imaging data should be equivalent to including age as a covariate to statistical adjustment by (de Lange et al., 2020), assuming optimal bias correction.

Given the established associations between neural oscillations/connectivity and age as measured by functional neuroimaging techniques such as EEG and MEG, and the ability to detect irregular/pathological activity, we hypothesise that functional neuroimaging derived measures of “brain age” are sensitive to atypical activity, such as that witnessed within neurodegenerative cohorts.

## Conclusion

Our findings suggest that strong associations exist between healthy ageing and neurophysiological features measured by MEG, such as spectral power and FC. In general, shifting spectral power from low to high frequencies across the whole brain, and decreasing intra and inter visual network connectivity were found to be correlated with healthy ageing. These associations were replicated using multiple independent techniques and were found to be spectrally and spatially consistent. Analysis of FC using MEG has previously been limited to cohorts composed of early childhood to midlife, here we expand into later life (18-83 years), extending the literature in this area. Cross-validated regression modelling was performed to predict participant age, with results showing mean absolute error of 8.54 years over mean intra-individual predictions.

## Author contributions

SH: Conceptualisation, Data curation, Formal analysis, Investigation, Methodology, Visualisation, Writing – original draft, Writing – review & editing.

GR: Conceptualisation, Data curation, Formal analysis, Investigation, Methodology, Visualisation, Writing – original draft, Writing – review & editing.

MV: Data curation, Investigation.

BTD: Conceptualisation, Investigation, Methodology, Project administration, Supervision, Writing – original draft, Writing – review & editing

## Acknowledgements

This work was funded by study operating sponsorship from MYndspan Ltd, disbursed to the lab of BTD, which supported data collection and partially the salary of MV.

## Appendix 1 Network nodes definitions

Central Executive Network

RIPS [25, -62, 53]

RVV [36, -62, 0]

LVV [-44, -60, -6]

RSMG [32, -38, 38]

RSLOC [26, -64, 54]

LSLOC [-26, -60, 52]

RFEF [28, -4, 58]

LFEF [-26, -8, 54]

Default Mode Network

LAG [-43, -76, 35]

RAG [51, -64, 32]

PCC [-3, -54, 31]

vMPFC [-2, 51, 2]

dMPFC [-13, 52, 23]

RMPFC [2, 53, 24]

LITG [-57, -25, -17]

Motor network

Precentral_L [-39.0, -7.0, 50.0]

Precentral_R [41.0, -10.0, 51.0]

Postcentral_L [-43.0, -24.0, 47.0]

Postcentral_R [41.0, -27.0, 51.0]

Parietal_Sup_L [-24.0, -61.0, 58.0]

Parietal_Sup_R [26.0, -60.0, 61.0]

Parietal_Inf_L [-43.0, -47.0, 45.0]

Parietal_Inf_R [46.0, -48.0, 48.0]

Attention network

RSMG [52, -48, 28]

RFEF [30, -13, 53]

LFEF [-26, -12, 53]

LpIPS [-25, -67, 48]

RpIPS [23, -69, 49]

LMT [-43, -72, -8]

RMT [42, -70, -11]

RMFG [41, 17, 31]

RPCS [41, 2, 50]

RSTG [58, -48, 10]

RVFC [40, 21, -4]

Visual network

LV1 [-3, -101, -1]

RV1 [11, -88, -4]

LV2d [-8, -99, 7]

RV2d [14, -96, 13]

LV3 [-9, -96, 13]

RV3 [20, -95, 18]

LV4 [-31, -77, -17]

RV4 [27, -71, -14]

LV7 [-23, -78, 26]

RV7 [32, -78, 25]

### Abbreviations

“RIPS”: “Right Intra Parietal Sulcus”
“RVV”: “Right Ventral Visual”
“LVV”: “Left Ventral Visual”
“RSMG”: “Right Supramarginal Gyrus”
“RSLOC”: “Right Superior Lateral Occipital Cortex”
“LSLOC”: “Left Superior Lateral Occipital Cortex”
“RFEF”: “Right Frontal Eye Field”
“LFEF”: “Left Frontal Eye Field”
“LAG”: “Left Angular Gyrus”
“RAG”: “Right Angular Gyrus”
“PCC”: “Posterior Cingulate Cortex”
“vMPFC”: “Ventromedial Prefrontal Cortex”
“dMPFC”: “Dorsomedial Prefrontal Cortex”
“RMPFC”: “Rostral Medial Prefrontal Cortex”
“LITG”: “Left Inferior Temporal Gyrus”
“Precentral_L”: “Precentral Left”
“Precentral_R”: “Precentral Right”
“Postcentral_L”: “Postcentral Left”
“Postcentral_R”: “Postcentral Right”
“Parietal_Sup_L”: “Parietal Superior Left”
“Parietal_Sup_R”: “Parietal Superior Right”
“Parietal_Inf_L”: “Parietal Inferior Left”
“Parietal_Inf_R”: “Parietal Inferior Right”
“LV1”: “Left Visual 1”
“RV1”: “Right Visual 1”
“LV2d”: “Left Visual 2 (Dorsal)”
“RV2d”: “Right Visual 2 (Dorsal)”
“LV3”: “Left Visual 3”
“RV3”: “Right Visual 3”
“LV4”: “Left Visual 4”
“RV4”: “Right Visual 4”
“LV7”: “Left Visual 7”
“RV7”: “Right Visual 7”
“LpIPS”: “Left posterior Intra-Parietal Sulcus”
“RpIPS”: “Right posterior Intra-Parietal Sulcus”
“LMT”: “Left Middle Temporal”
“RMT”: “Right Middle Temporal”
“RMFG”: “Right Middle Frontal Gyrus”
“RPCS”: “Right Precentral Sulcus”
“RSTG”: “Right Superior Temporal Gyrus”
“RVFC”: “Right Ventro-Frontal Cortex”

## Appendix 2 Partial Correlation

To assess the trends of spectral and connectivity features with respect to age we used partial correlation, which allows us to quantify the relationship between two continuous variables while removing the effects of any potentially confounding variables. In practice this allows us to confidently assess the effect of age on functional activity, while removing the effects of sex. In the case of multiple confounding variables, the partial correlation coefficient is calculated by first performing a multiple linear regression in which confounding variables are considered regressors alongside one of the continuous variables of the correlation, the second of the continuous variables is set as the response variable. Once regression has been performed, equation 1 is used to transform the regression results for a continuous variable into the partial correlation coefficient, where *t_f_* is the student’s t-test of the multiple regression coefficient *β_f_*, *p* is the number of explanatory variables, and *n* is the number of samples.

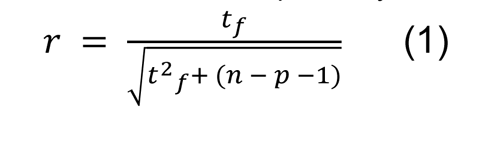

Tests are performed by using one MEG derived feature, chronological age, and all relevant confounding variables (sex). Each of these tests was replicated on 1000 bootstrapped samples such that confidence intervals of the correlation coefficients can be calculated. Permutation tests were used to further assess the statistical significance of the tests, by generating a null distribution of the test statistic to calculate a p-value via comparison to the test statistic found using the full population.

The results of the partial correlation analysis show a general trend of oscillatory slowing associated with age, as delta and theta band power decreases, and beta and gamma activity increases. Spectral power across bands was generally more highly correlated and statistically significant than connectivity, which resulted in a higher proportion of the Bonferroni corrected results originating from the spectral features.

The results of the partial correlation analysis of spectral power show age dependent features of healthy ageing which are temporally and regionally specific, as well as highly bilateral. Delta power decreases across all but the occipital lobe with ageing (**Figure S1a**), similarly theta activity tends to decrease, with temporal and parietal regions presenting the strongest association with age (**Figure S1b**). Alpha spectral activity has the weakest associations with age, and is the only frequency band with bidirectional changes, with a subset of occipital and parietal regions showing decreasing power, and conversely temporal and frontal areas showing increasing activity (**Figure S1c**). Power in the beta band displays the strongest correlations with age (**Figure S1d**), with all regions proving statistically significant post multiple comparisons correction. Gamma spectral power continues the trend of oscillatory slowing, as positive correlations are present primarily in the occipital and frontal lobes (**Figure S1e**).

Across the canonical frequency bands of delta, theta, alpha, beta, and gamma, the left middle frontal gyrus orbital part (***r***(367)=-.56, ***p***=2.81e-24), right precentral (***r***(367)=-.56, ***p***=1.27e-22), right cuneus (***r***(367)=-.36, ***p***=3.84e-9), left postcentral (***r***(367)=.64, ***p***=2.68e-29), and right lingual (***r***(367)=.48, ***p***=7.26e-18) regions had the greatest age association respectively.

**Figure S1:**
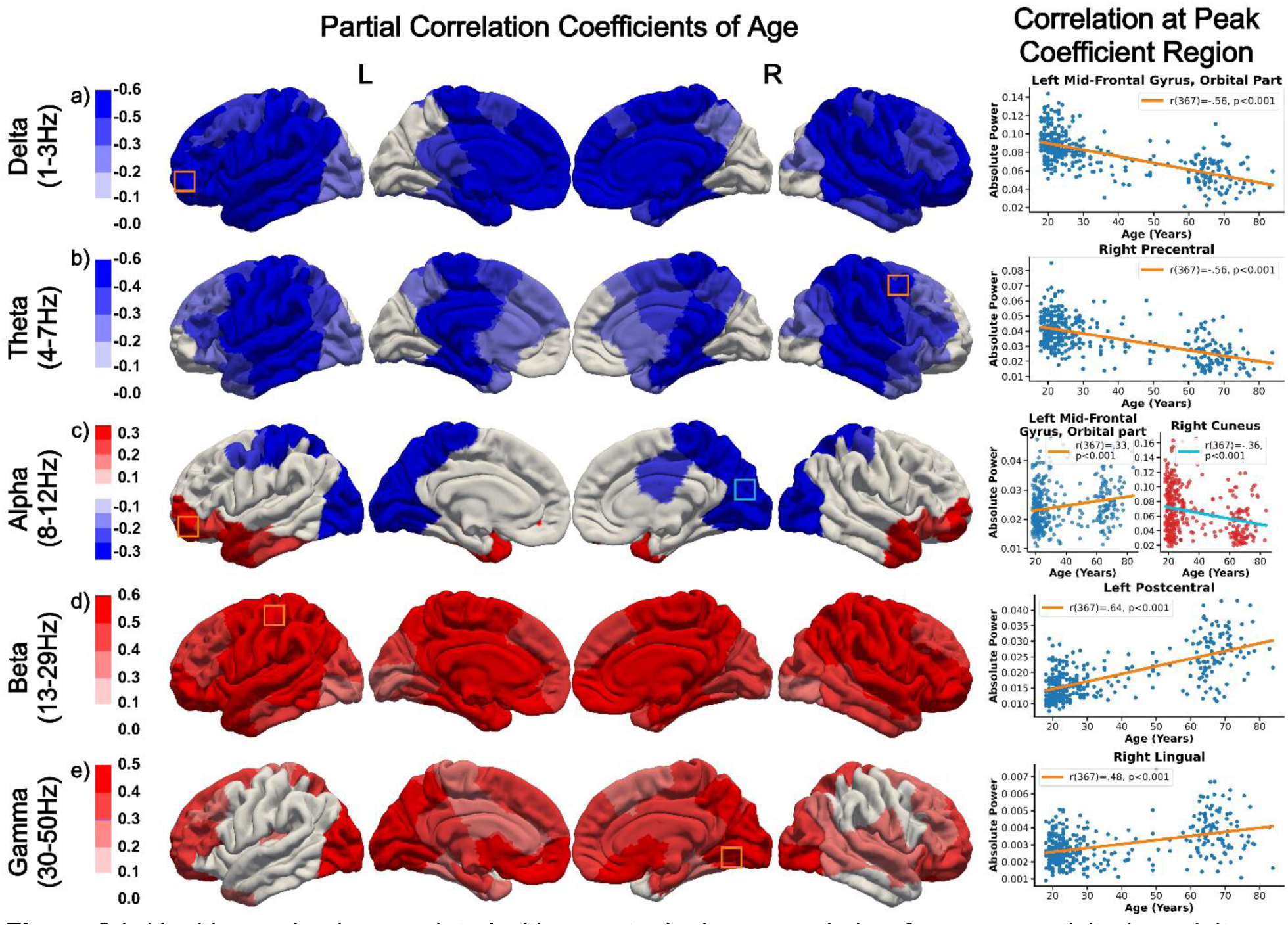
Healthy ageing is associated with monotonic decreases in low frequency activity (e.g. delta and theta), and simultaneous increases in high frequency activity (e.g. beta and gamma), with regionally dependent bidirectional changes in alpha across prefrontal, temporal, and occipital locations. The strongest relationships between healthy ageing and spectral power were found in beta, which presents widespread increases across the brain to be associated with healthy ageing. These results show the Bonferroni corrected partial correlation coefficients of spectral power and healthy ageing. Bounding boxes on the band limited power plots highlight partial correlation coefficient peak regions, the colour of the bounding box corresponds to the line of best fit shown on the adjacent scatter plot, which shows data from this region.

Decreased connectivity in the theta band was associated with age in a limited number of inter-network connections of the central executive, motor, and attention networks. Alpha connectivity likewise shows decreases with age, primarily in the visual network intra and inter network connections. Beta connectivity increased in a subset of default mode intra-network connections, and conversely beta and gamma connectivity decreased in subsets of visual intra-network connections.

**Figure S2:**
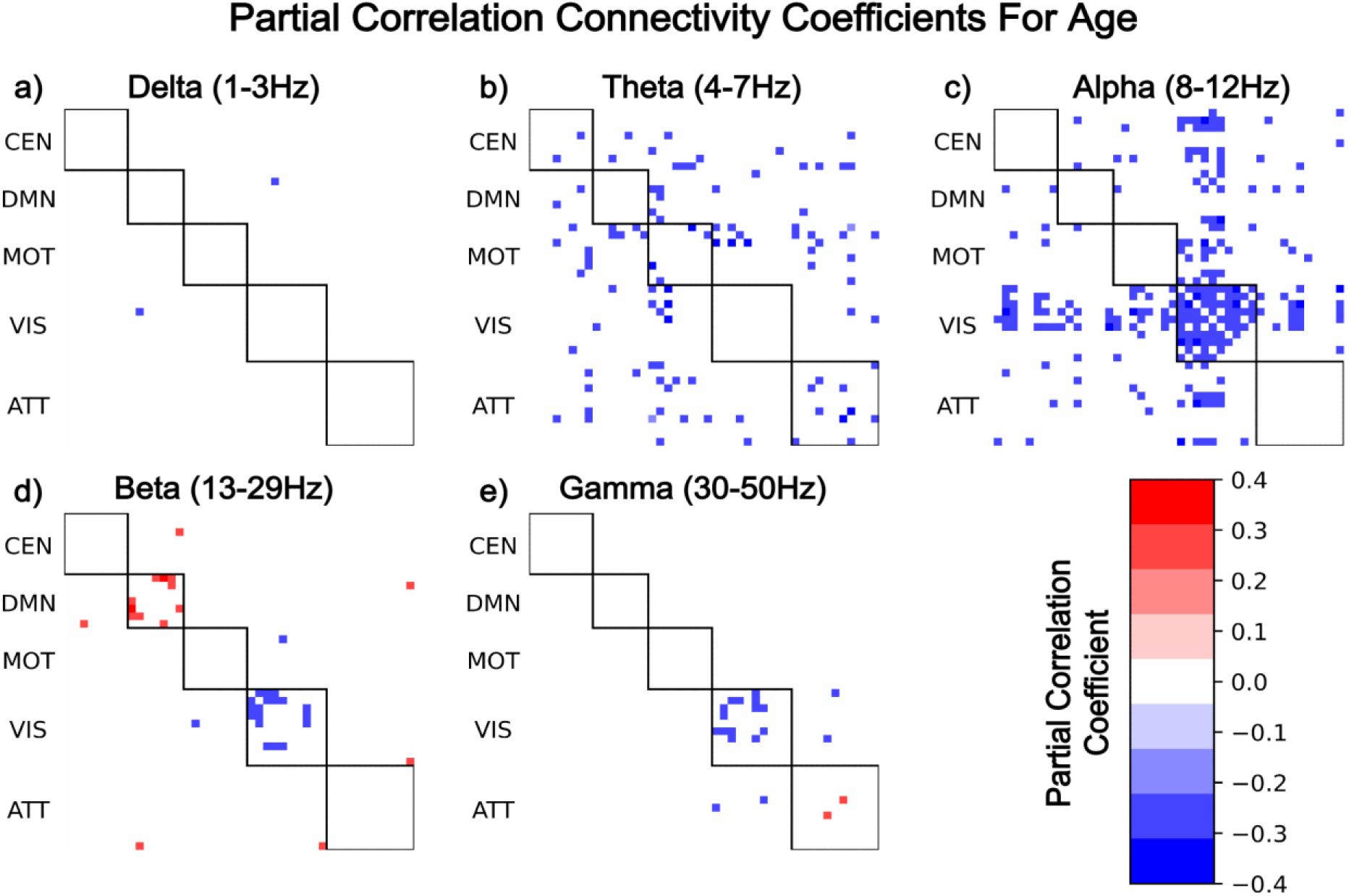
Healthy ageing is associated with decreasing intra visual network connectivity in alpha, beta, and gamma, as well as decreasing inter visual network connectivity in alpha. Increasing default mode network connectivity in the beta band was found in a subset of connections. These results show the Bonferroni corrected partial correlation coefficients of functional connectivity and healthy ageing.

To establish the degree of coherence between the PLS weights and partial correlation coefficients of the spectral power and FC features we used spearman’s rank correlation. The results of this analysis are shown in **Table S1** and highlight that although associations between age and spectral power features were on average more consistent across different techniques than FC, the relationships between beta AEC and age were also highly stable. The lower correlation between PLS weights and partial correlation coefficients for AEC is potentially motivated by the lack of a clear age association in many inter-network connections. Alternatively, lack of correlation between these two measures could be the result of non-linear relationships between age and a given feature or set of features, as spearman’s rank correlation used in partial correlation captures monotonic relationships rather than the linear association of PLS when considered at the non-latent variable level.

**Table S1:**
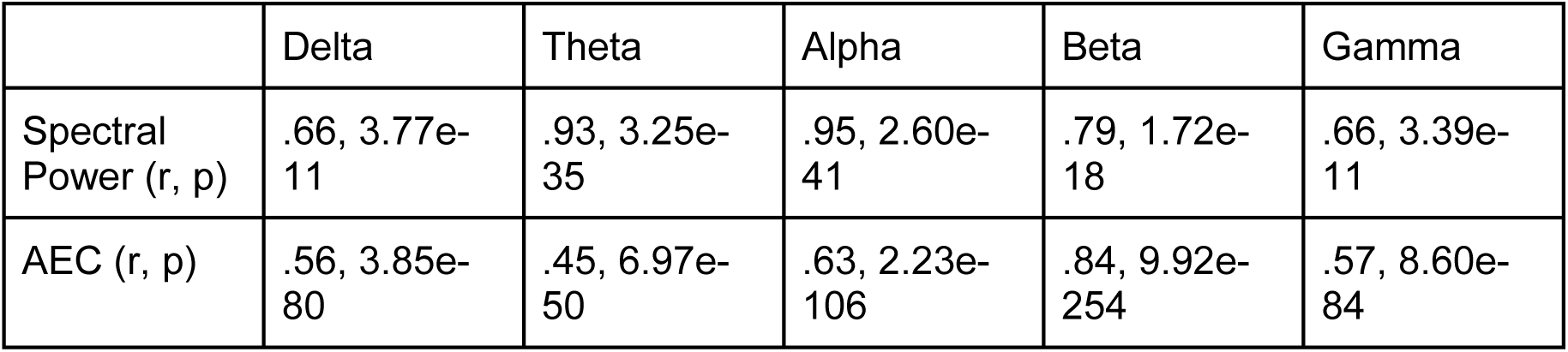
Correlation between PLS weights and partial correlation coefficients demonstrates the degree of coherence in the derived effects of healthy ageing by the two methods. Alpha and theta spectral power show the greatest correlation between methods, followed by beta functional connectivity, providing credence to the notion of beta connectivity being the most stable and implicated frequency band for connectivity throughout the brain.

