## Appendix for "Predicting brain age across the adult lifespan with spontaneous oscillations and functional coupling in resting brain networks captured with magnetoencephalography"

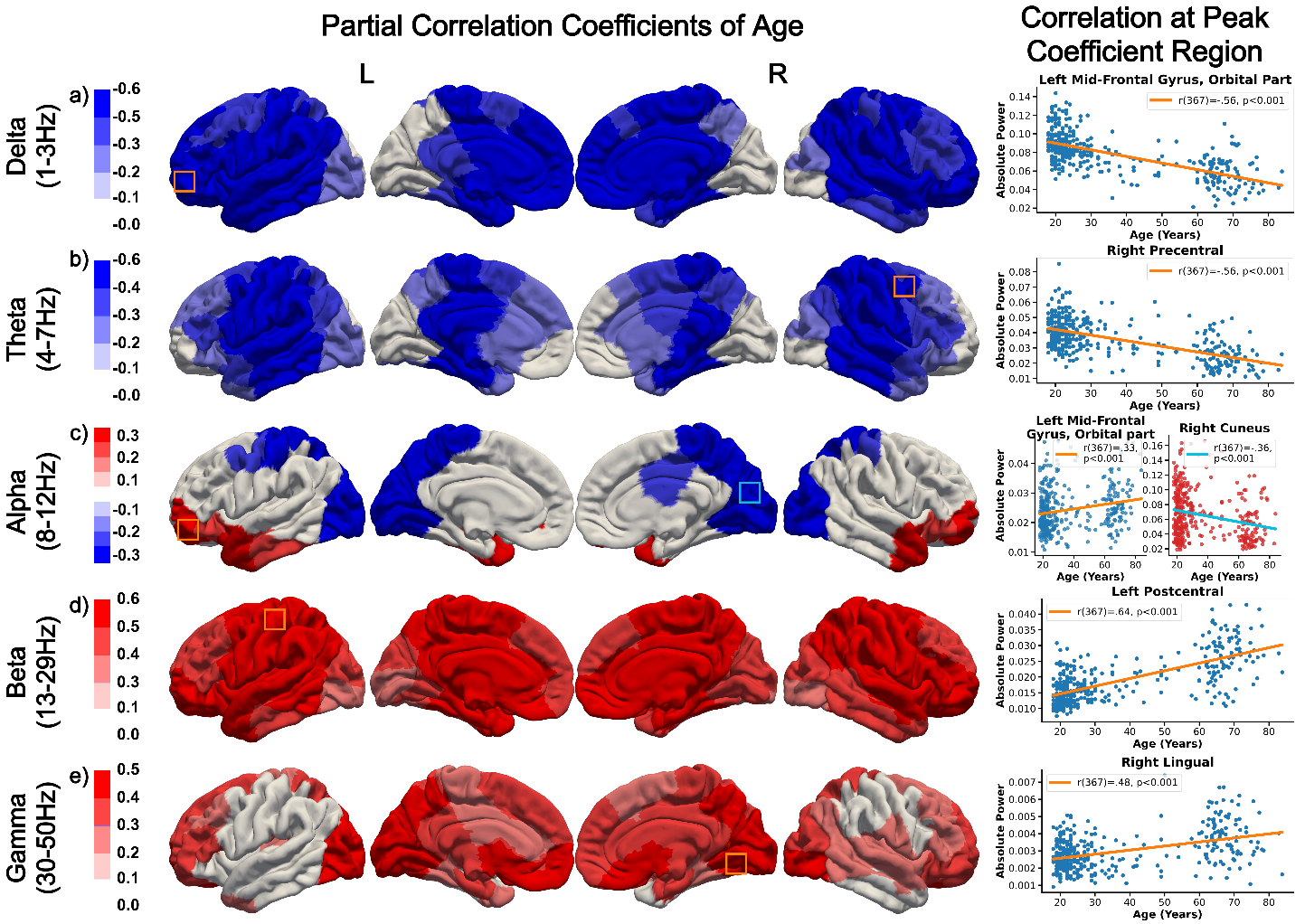
***Figure S1:*** *Healthy ageing is associated with monotonic decreases in low frequency activity (e.g. delta and theta), and simultaneous increases in high frequency activity (e.g. beta and gamma), with regionally dependent bidirectional changes in alpha across prefrontal, temporal, and occipital locations. The strongest relationships between healthy ageing and spectral power were found in beta, which presents widespread increases across the brain to be associated with healthy ageing. These results show the Bonferroni corrected partial correlation coefficients of spectral power and healthy ageing. Bounding boxes on the band limited power plots highlight partial correlation coefficient peak regions, the colour of the bounding box corresponds to the line of best fit shown on the adjacent scatter plot, which shows data from this region.*

Decreased connectivity in the theta band was associated with age in a limited number of inter-network connections of the central executive, motor, and attention networks. Alpha connectivity likewise shows decreases with age, primarily in the visual network intra and inter network connections. Beta connectivity increased in a subset of default mode intra-network connections, and conversely beta and gamma connectivity decreased in subsets of visual intra-network connections.

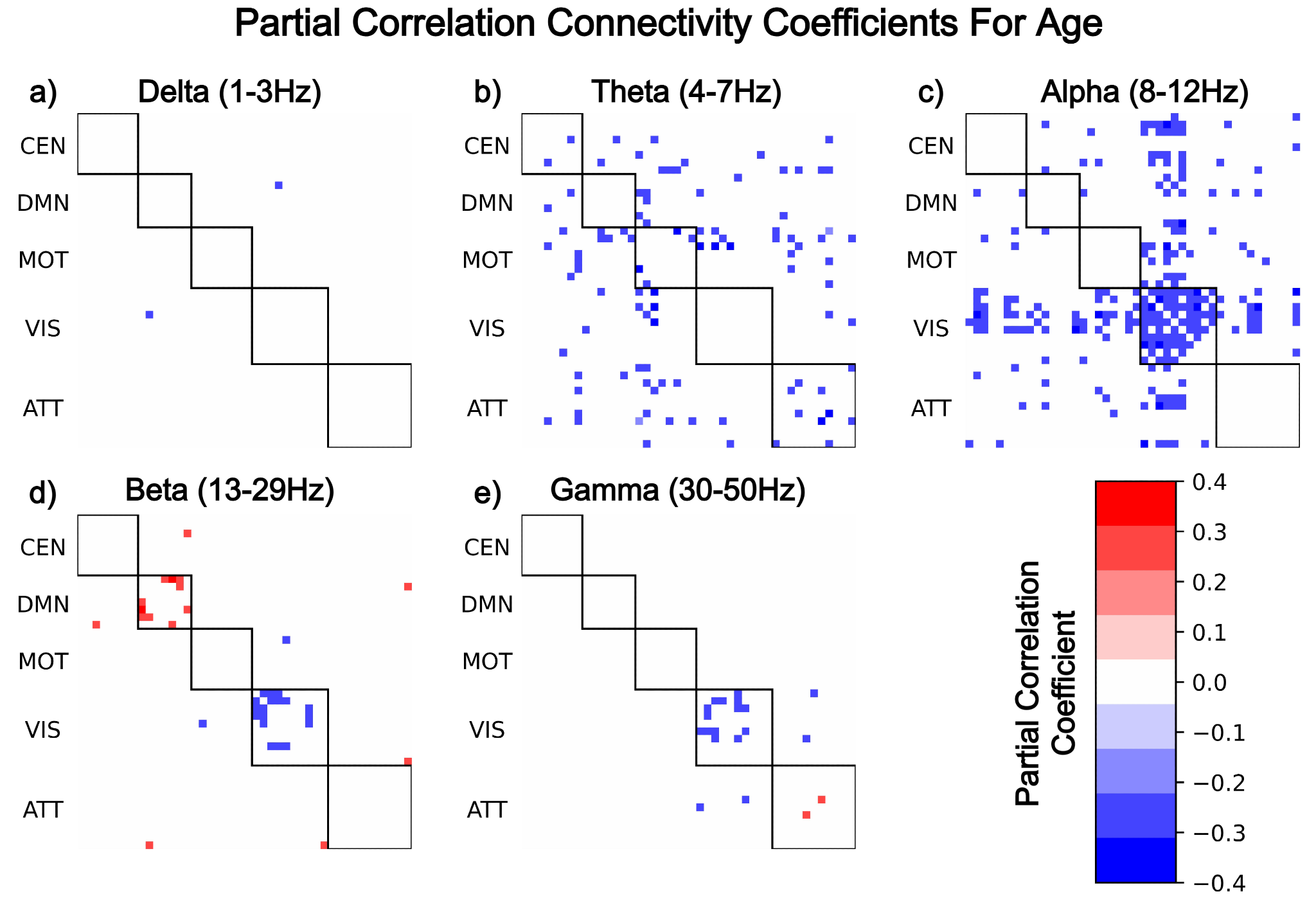

***Figure S2:*** *Healthy ageing is associated with decreasing intra visual network connectivity in alpha, beta, and gamma, as well as decreasing inter visual network connectivity in alpha. Increasing default mode network connectivity in the beta band was found in a subset of connections. These results show the Bonferroni corrected partial correlation coefficients of functional connectivity and healthy ageing.*

To establish the degree of coherence between the PLS weights and partial correlation coefficients of the spectral power and FC features we used spearman's rank correlation. The results of this analysis are shown in **Table S1** and highlight that although associations between age and spectral power features were on average more consistent across different techniques than FC, the relationships between beta AEC and age were also highly stable. The lower correlation between PLS weights and partial correlation coefficients for AEC is potentially motivated by the lack of a clear age association in many inter-network connections. Alternatively, lack of correlation between these two measures could be the result of non-linear relationships between age and a given feature or set of features, as spearman’s rank correlation used in partial correlation captures monotonic relationships rather than the linear association of PLS when considered at the non-latent variable level.

***Table S1:*** *Correlation between PLS weights and partial correlation coefficients demonstrates the degree of coherence in the derived effects of healthy ageing by the two methods. Alpha and theta spectral power show the greatest correlation between methods, followed by beta functional connectivity, providing credence to the notion of beta connectivity being the most stable and implicated frequency band for connectivity throughout the brain.*

|  | Delta | Theta | Alpha | Beta | Gamma |
| --- | --- | --- | --- | --- | --- |
| Spectral Power (r, p) | .66, 3.77e-11 | .93, 3.25e-35 | .95, 2.60e-41 | .79, 1.72e-18 | .66, 3.39e-11 |
| AEC (r, p) | .56, 3.85e-80 | .45, 6.97e-50 | .63, 2.23e-106 | .84, 9.92e-254 | .57, 8.60e-84 |
